# Manganese is a Physiologically Relevant TORC1 Activator in Yeast and Mammals

**DOI:** 10.1101/2021.12.09.471923

**Authors:** Raffaele Nicastro, Hélène Gaillard, Laura Zarzuela, Marie-Pierre Péli-Gulli, Elisabet Fernández-García, Mercedes Tomé, Néstor García-Rodríguez, Raúl V. Dúran, Claudio De Virgilio, Ralf Erik Wellinger

## Abstract

The essential biometal manganese (Mn) serves as a cofactor for several enzymes that are crucial for the prevention of human diseases. Whether intracellular Mn levels may be sensed and modulate intracellular signaling events has so far remained largely unexplored. The highly conserved target of rapamycin complex 1 (TORC1, mTORC1 in mammals) protein kinase requires divalent metal cofactors such as magnesium (Mg^2+^) to phosphorylate effectors as part of a homeostatic process that coordinates cell growth and metabolism with nutrient and/or growth factor availability. Here, our genetic approaches reveal that TORC1 activity is stimulated *in vivo* by elevated cytoplasmic Mn levels, which can be induced by loss of the Golgi-resident Mn^2+^ transporter Pmr1 and which depends on the natural resistance-associated macrophage protein (NRAMP) metal ion transporters Smf1 and Smf2. Accordingly, genetic interventions that increase cytoplasmic Mn^2+^ levels antagonize the effects of rapamycin in triggering autophagy, mitophagy, and Rtg1-Rtg3-dependent mitochondrion-to-nucleus retrograde signaling. Surprisingly, our *in vitro* protein kinase assays uncovered that Mn^2+^ activates TORC1 substantially better than Mg^2+^, which is primarily due to its ability to lower the K_m_ for ATP, thereby allowing more efficient ATP coordination in the catalytic cleft of TORC1. These findings, therefore, provide both a mechanism to explain our genetic observations in yeast and a rationale for how fluctuations in trace amounts of Mn can become physiologically relevant. Supporting this notion, TORC1 is also wired to feedback control mechanisms that impinge on Smf1 and Smf2. Finally, we also show that Mn^2+^-mediated control of TORC1 is evolutionarily conserved in mammals, which may prove relevant for our understanding of the role of Mn in human diseases.

**Significance Statement:** The target of rapamycin complex 1 (TORC1, mTORC1 in mammals) is a central, highly conserved controller of cell growth and aging in eukaryotes. Our study shows that the essential biometal manganese (Mn) acts as a primordial activator of TORC1 and that NRAMP metal ion transporters control TORC1 activity by regulating cytoplasmic Mn^2+^ levels. Moreover, TORC1 activity regulates Mn^2+^ levels through feedback circuits impinging on NRAMP transporters. Altogether, our results indicate that Mn homeostasis is highly regulated and modulates key cellular processes such as autophagy, mitophagy, and Rtg1-3 complex-dependent retrograde response. These findings open new perspectives for the understanding of neurodegenerative disorders and aging-related processes

## Introduction

Mn is a vital trace element that is required for the normal activity of the brain and nervous system by acting, among other mechanisms, as an essential, divalent metal cofactor for enzymes such as the mitochondrial enzyme superoxide dismutase 2 (1), the apical activator of the DNA damage response serine/threonine kinase ATM (2) or the Mn^2+^-activated glutamine synthetase (3). However, Mn^2+^ becomes toxic when enriched in the human body (4). While mitochondria have been proposed as a preferential organelle where Mn^2+^ accumulates and unfolds its toxicity by increasing oxidative stress and thus mitochondrial dysfunction (5), the molecular mechanisms of Mn^2+^ toxicity in humans are also related to protein misfolding, endoplasmic reticulum (ER) stress, and apoptosis (6). Mn^2+^ homeostasis is coordinated by a complex interplay between various metal transporters for Mn^2+^ uptake and intracellular Mn^2+^ distribution and represents an essential task of eukaryotic cells, which is also of vital importance specifically for neuronal cell health (7).

Much of our knowledge on Mn^2+^ transport across the plasma membrane into the ER, the Golgi, endosomes, and vacuoles comes from studies in *Saccharomyces cerevisiae* (outlined in Figure 1A). Typically, Mn^2+^ is shuttled across membranes by transporters that belong to the NRAMP family, which are highly conserved metal transporters responsible for iron (Fe) and Mn^2+^ uptake (8). Not surprisingly, therefore, NRAMP orthologs have been found to cross-complement functions in yeast, mice, and humans (9). One of the best-studied NRAMPs is the yeast plasma membrane protein Smf1. Interestingly, extracellular Fe or Mn^2+^ supplementation triggers Bsd2 adaptor protein-dependent, Rsp5-mediated ubiquitination of Smf1, which initiates its sorting through the endocytic multivesicular body pathway and subsequent lysosomal degradation (10, 11). The Smf1 paralogs Smf2 and Smf3 are less well studied, but Smf2 is predominantly localized at endosomes and its levels decrease under conditions of Mn or Fe overload (12, 13). Within cells, the P-type ATPase Pmr1 (also known as Bsd1) represents a key transporter that shuttles Ca^2+^ and Mn^2+^ ions into the Golgi lumen. Its loss leads to increased levels of Mn^2+^ in the cytoplasm due to defective detoxification (14). Noteworthy, several phenotypes associated with loss of Pmr1 have been shown to arise as a consequence of Mn^2+^ accumulation in the cytoplasm, including telomere shortening, genome instability and bypass of the superoxide dismutase Sod1 requirement (14–17).

**Figure 1.**
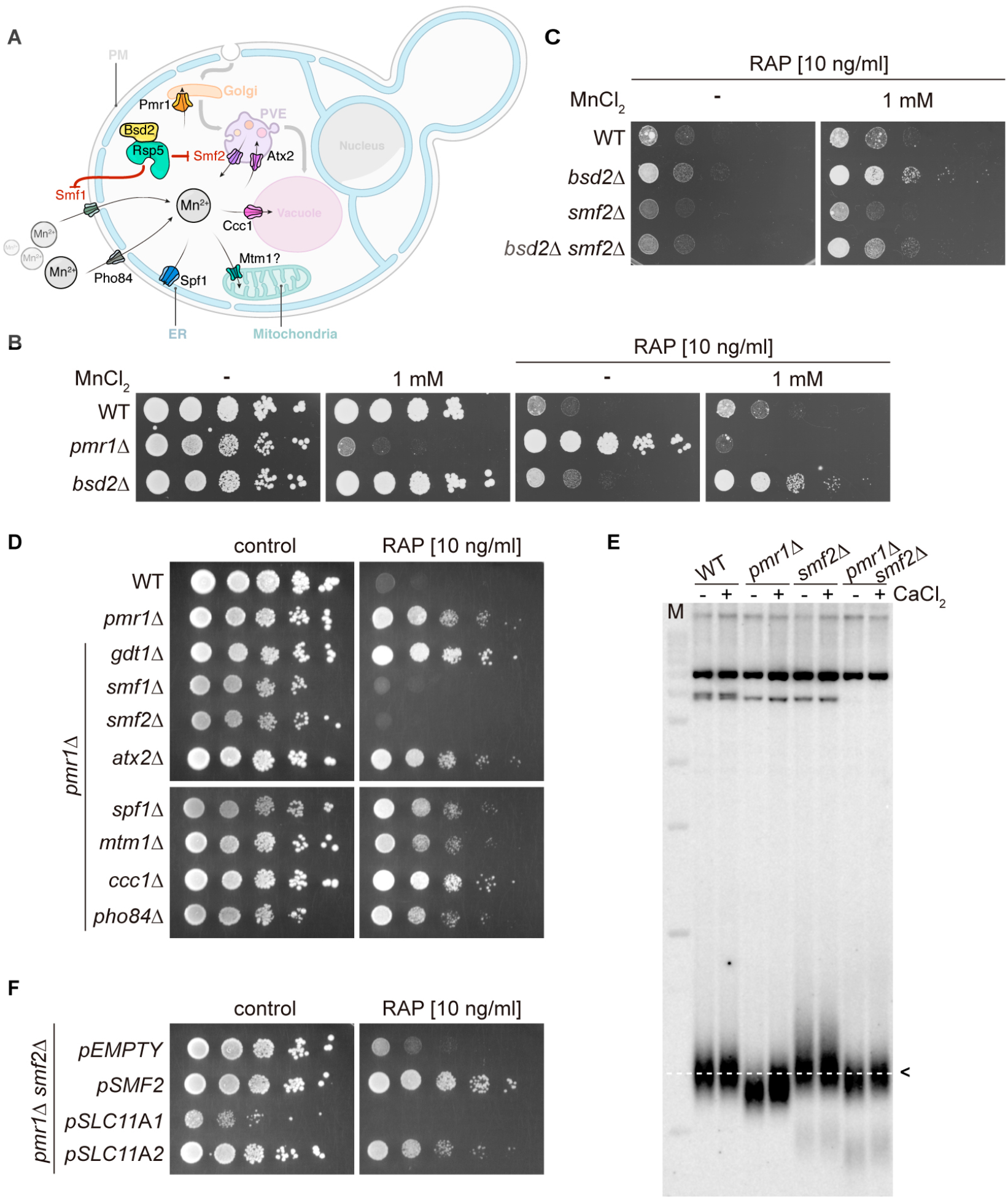
NRAMP transporters link Mn-import to rapamycin resistance. (**A**) Schematical outline of yeast Mn^2+^-transporters and their intracellular localization. PM, plasma membrane; PVE, pre-vacuolar endosomes; ER, endoplasmic reticulum. Note that Bsd2 is a specific adaptor protein for Rsp5-mediated Smf1 and Smf2 ubiquitination in response to Mn^2+^ overload. The Golgi Mn^2+^-transporter Gdt1 is omitted for clarity. (**B-D**) Growth on MnCl_2_- and/or rapamycin-containing medium (RAP). 10-fold dilutions of exponentially growing cells are shown. Strains and compound concentrations are indicated. Note that the medium used in (D) was supplemented with 10 mM CaCl_2_. Data obtained in a medium without CaCl_2_ are shown in Figure S3. (**E**) Southern blot analysis of telomere length. Genomic DNA was derived from cells grown in a medium supplemented or not with CaCl_2_ and cleaved by *Xho*I before agarose gel electrophoresis. The 1.3 kb average length of telomeres from WT cells (dashed white line, black arrow) and size marker (M) are shown. (**F**) Growth of *pmr1*Δ *smf2*Δ double mutants transformed with plasmids expressing yeast Smf2, *Mus musculus* SLC11A1, or SLC11A2 on rapamycin-containing medium.

TORC1/mTORC1 is a central, highly conserved controller of cell growth and aging in eukaryotes. It coordinates the cellular response to multiple inputs, including nutritional availability, bioenergetic status, oxygen levels, and, in multicellular organisms, the presence of growth factors (18, 19). In response to these diverse cues, TORC1 regulates cell growth and proliferation, metabolism, protein synthesis, autophagy, and DNA damage responses (20, 21). In *S. cerevisiae*, which played a pivotal role in the discovery and dissection of the TOR signaling network (22), TORC1 is mainly localized on the surfaces of vacuoles and endosomes (23, 24) where it integrates, among other cues, amino acid signals through the Rag GTPases and Pib2 (25, 26). In mammals, amino acids also activate the Rag GTPases, which then recruit mTORC1 to the lysosomal surface where it can be allosterically activated by the small GTPase RHEB (Ras homolog expressed in brain) that mediates the presence of growth factors and sufficient energy levels (20, 27–29). Interestingly, mTORC1 regulates cellular Fe homeostasis (30), but how it may be able to sense Fe levels remains largely unknown. In addition, although TORC1/mTORC1 requires divalent metal ions to coordinate ATP at its catalytic cleft (31, 32), it is currently not known whether these or any other trace elements may play a physiological or regulatory role in controlling its activity.

In yeast, genetic evidence links high levels of cytoplasmic Mn^2+^ levels to TORC1 function, as loss of Pmr1 confers rapamycin resistance (33). However, the underlying molecular mechanism(s) by which Mn^2+^ may mediate rapamycin resistance remains to be explored. Here, we show that Mn^2+^ uptake by NRAMP transporters modulates rapamycin resistance and that, in turn, TORC1 inhibition by rapamycin regulates NRAMP transporter availability. Moreover, intracellular Mn^2+^ excess antagonizes rapamycin-induced autophagy, mitophagy, and Rtg1-3 transcription factor complex-dependent retrograde response activation. Surprisingly, our *in vitro* and *in vivo* analyses reveal that TORC1 protein kinase activity is strongly activated in the presence of MnCl_2_. In our attempts to understand the mechanisms underlying these observations, we discovered that Mn^2+^, when compared to Mg^2+^, significantly boosts the affinity of TORC1 for ATP. Combined, our findings also indicate that TORC1 activity is regulated by and regulates intracellular Mn^2+^ levels, defining Mn^2+^ homeostasis as a key factor in cell growth control. Importantly, our studies in human cells indicate that Mn^2+^-driven TORC1 activation is likely conserved throughout evolution, opening new perspectives for our understanding of Mn^2+^ toxicities and their role in neurodegenerative disorders and aging.

## Results

### NRAMP transporters regulate cytoplasmic Mn^2+^ levels and rapamycin resistance

Yeast cells lacking the Golgi-localized P-type ATPase Pmr1, which transports Ca^2+^ and Mn^2+^ ions from the cytoplasm to the Golgi lumen, are resistant to the TORC1 inhibitor rapamycin (33, 34), a phenotype that is generally associated with increased TORC1 activity. In *pmr1*Δ mutants, defective Mn^2+^ shuttling at the Golgi leads to protein sorting defects and accumulation of the general amino-acid permease Gap1 at the plasma membrane (35). In theory, this may translate into unrestrained uptake and intracellular accumulation of amino acids, and thus hyperactivation of TORC1. However, arguing against such a model, we found cells lacking both Pmr1 and Gap1 to remain resistant to low doses of rapamycin (Figure S1A). We then asked whether loss of Pmr1 may affect the expression levels or cellular localization of TORC1. Our results indicated that GFP-Tor1 protein levels and localization to vacuolar and endosomal membranes remained unaltered in exponentially growing and rapamycin-treated WT and *pmr1*Δ cells (Figure S1B and S1C). Given the roughly 5-fold increased intracellular Mn^2+^ levels of cells lacking Pmr1 (14), we then considered the possibility that Mn^2+^ may have a more direct role in TORC1 activation. We, therefore, assessed the rapamycin sensitivity of cells lacking the adaptor protein Bsd2, which mediates Rsp5-dependent degradation in response to high Mn^2+^ levels of both the plasma membrane- and endosomal membrane-resident NRAMP Mn^2+^ transporters Smf1 and Smf2, respectively (Figure 1A; (12)). Interestingly, *bsd2*Δ cells were as sensitive to rapamycin as WT cells, but, unlike WT cells, could be rendered rapamycin resistant by the addition of 1 mM MnCl_2_ in the growth medium (Figure 1B and Figure S2). This effect was strongly reduced in the absence of Smf2 (Figure 1C), suggesting that Smf2-dependent endosomal Mn^2+^ export and, consequently, cytoplasmic accumulation of Mn^2+^ may be required for rapamycin resistance under these conditions. In line with such a model, we found that overexpression of the vacuolar membrane-resident Vcx1-M1 transporter, which imports Mn^2+^ into the vacuolar lumen, suppressed the Mn-induced rapamycin resistance of *bsd2*Δ mutants and increased their sensitivity to rapamycin in the absence of extracellular MnCl_2_ supply (Figure S2).

As schematically outlined in Figure 1A, several metal transporters have been associated with Mn^2+^transport in yeast, including those localized at the plasma membrane (Smf1 and Pho84), the endosomes (Smf2 and Atx2), the Golgi (Pmr1 and Gdt1), the vacuole (Ccc1 and Vcx1), the ER (Spf1), and possibly the mitochondria (Mtm1). To identify which of these transporters contributes to the rapamycin resistance of *pmr1*Δ cells, we next monitored the growth of double mutants lacking Pmr1 and any of the corresponding metal transporters in the presence of rapamycin (Figure 1D). Interestingly, only loss of Smf1 or Smf2 reestablished rapamycin sensitivity in *pmr1*Δ cells, while loss of Gdt1 or Ccc1 even slightly enhanced the rapamycin-resistance phenotype of *pmr1*Δ cells. Of note, growth was monitored on CaCl_2_-supplemented media that improved the growth of *pmr1*Δ cells lacking Smf1 or Smf2 without affecting their sensitivity to rapamycin, which also rules out the possibility that Ca^2+^ is associated with the observed rapamycin resistance phenotype (Figure 1D and Figure S3). Taken together, our data indicate that increased intracellular Mn^2+^ levels in *pmr1*Δ cells lead to rapamycin resistance and that this phenotype can be suppressed either by reducing Mn^2+^ import through loss of Smf1 or, possibly, by increasing Mn^2+^sequestration in endosomes through loss of Smf2 as previously suggested (36). To confirm our prediction that loss of Smf2 reduces cytoplasmic Mn^2+^ levels, we measured MnCl_2_-dependent telomere length shortening as an indirect proxy for cytoplasmic and nuclear Mn levels (37). Accordingly, telomere length was significantly decreased in *pmr1*Δ cells (when compared to WT cells), increased in *smf2*Δ *cells*, and similar between *pmr1*Δ *smf2*Δ and WT cells (Figure 1E). These data, therefore, corroborate our assumption that loss of Smf2 suppresses the high cytoplasmic and nuclear Mn^2+^ levels of *pmr1*Δ cells.

The function of metal transporters is highly conserved across evolution as exemplified by the fact that the expression of the human Pmr1 ortholog, the secretory pathway Ca^2+^/Mn^2+^ ATPase ATP2C1/SPCA1 (ATPase secretory pathway Ca^2+^ transporting 1), can substitute for Pmr1 function in yeast (38). A similar degree of functional conservation from lower to higher eukaryotes exists for NRAMP transporters (9). Because Smf2 is of specific interest in the context of the present study, we asked whether the orthologous mouse proteins, *i.e.* the divalent metal transporter SLC11A1 (NRAMP1) and SLC11A2 (DMT1) isoforms (solute carrier family 11 member 1 and 2, respectively), can restore rapamycin resistance in *pmr1*Δ *smf2*Δ double mutants. This was indeed the case, as the SLC11A2 isoform complemented Smf2 function in these assays (Figure 1F), indicating that Pmr1 and Smf2 are evolutionarily conserved transporters that are required for Mn^2+^ homeostasis.

### Elevated levels of intracellular Mn^2+^ antagonize rapamycin-induced autophagy, mitophagy, and Rtg1-3 retrograde signaling

Growth inhibition by rapamycin mimics starvation conditions and leads to the degradation and recycling of a wide spectrum of biological macromolecules via autophagy. In this context, the *pmr1*Δ mutant has previously been found to be defective in nutrient depletion-induced mitophagy (39). We thus wondered if rapamycin-induced autophagic processing may also be defective in *pmr1*Δ mutants. We took advantage of a GFP-Atg8 fusion construct (40) to monitor autophagy through GFP-Atg8 synthesis and processing in cells that had been subjected to rapamycin treatment for up to 6 hours. Rapamycin-induced GFP-Atg8 expression was strongly reduced in *pmr1*Δ cells and this phenotype was mitigated in the *pmr1*Δ *smf2*Δ double mutant (Figure 2A and B), suggesting that the initiation of autophagy is compromised by elevated cytoplasmic Mn^2+^levels. Next, we assessed mitophagy by following the expression and degradation of the GFP-tagged mitochondrial membrane protein Om45 (41). Rapamycin treatment led to a time-dependent up-regulation Om45-GFP protein levels in all tested strains (Figure 2C and D). However, the accumulation of the cleaved GFP protein was strongly reduced in the *pmr1*Δ single mutant, while this effect was again suppressed in the *pmr1*Δ *smf2*Δ double mutant. Combined, our results therefore suggest that the intracellular Mn^2+^ flux modulates both rapamycin-induced autophagy and mitophagy.

**Figure 2.**
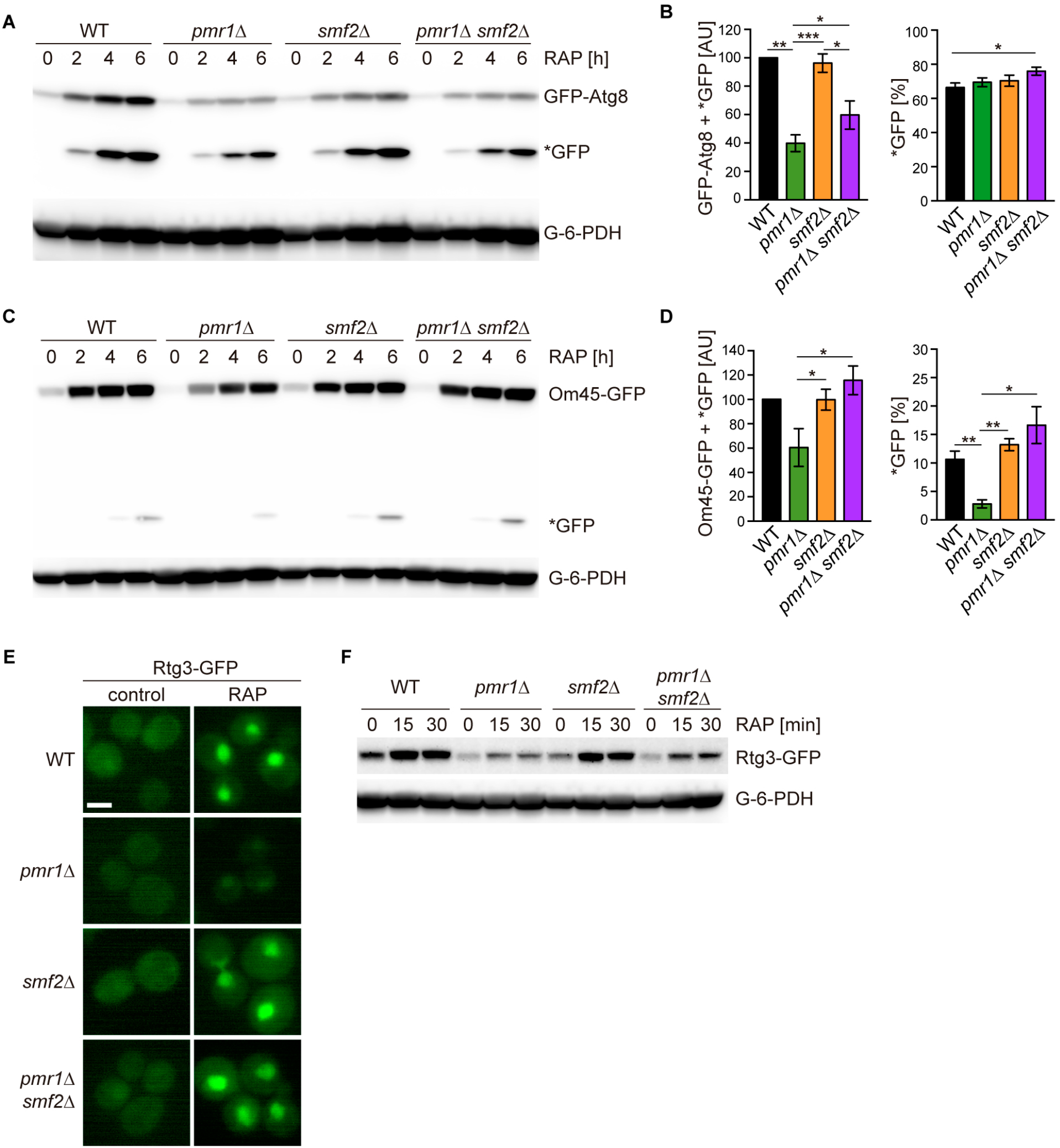
Intracellular Mn excess antagonizes rapamycin-induced autophagy, mitophagy, and Rtg1-3 retrograde signaling. (**A**) Exponentially growing WT and indicated mutant strains expressing plasmid-encoded GFP-Atg8 were treated for up to 6 h with 200 ng/ml rapamycin (RAP). GFP-Atg8 and cleaved GFP (*GFP) protein levels were analyzed by immunoblotting. Glucose-6-phosphate dehydrogenase (G-6-PDH) levels were used as a loading control. (**B**) Quantification of GFP-Atg8 and *GFP levels after a 6 h rapamycin treatment. Total GFP signal (Atg8-GFP + *GFP) normalized to WT levels (left) and percentage of *GFP relative to the total GFP signal (right) are plotted. Data represent means ± SEM of independent experiments (*n*=4). Statistical analysis: two-tailed t-test (paired for normalized data, unpaired for GFP* percentage). * p<0.05; ** p<0.01; *** p<0.001. (**C**) Exponentially growing WT and indicated mutant strains expressing Om45-GFP from the endogenous locus were treated and processed as in (A). (**D**) Quantification of Om45-GFP and *GFP levels after a 6 h rapamycin treatment. Details as in (B) with n=3. (**E**) Representative fluorescence microscope images of WT, *pmr1*Δ, *smf2*Δ, and *pmr1*Δ *smf2*Δ mutants expressing an episomic Rtg3-GFP reporter construct. Exponentially growing cells were treated or not (control) for 30 min with 200 ng/ml rapamycin (RAP). Scale bar represents 5 μm. (**F**) Rtg3-GFP expressing cells were treated for up to 30 min with 200 ng/ml rapamycin. protein levels were analyzed by immunoblotting. G-6-PDH levels were used as a loading control.

To identify additional responses to elevated cytosolic Mn^2+^ levels, we took advantage of our previously published transcriptome analysis of *pmr1*Δ cells (37). Careful analysis of these datasets revealed that the expression levels of genes activated by the heterodimeric Rtg1-Rtg3 transcription factor are reduced in *pmr1*Δ cells (see Table S1). Since TORC1 inhibits cytoplasmic-to-nuclear translocation of Rtg1-Rtg3 (42), we monitored the localization and protein levels of Rtg3-GFP in exponentially growing and rapamycin-treated WT and *pmr1*Δ cells. Rapamycin treatment not only induced nuclear enrichment of Rtg3-GFP as reported (Figure 2E; (42)) but also significantly increased the levels of Rtg3-GFP (Figure 2F). Loss of Pmr1, in contrast, significantly reduced the Rtg3-GFP levels in exponentially growing and rapamycin-treated cells, which also translated into barely visible levels of Rtg3-GFP in the nucleus (Figure 2E and 2F). Importantly, and in line with our finding that loss of Smf2 suppresses the high cytoplasmic and nuclear Mn^2+^ levels of *pmr1*Δ cells (see above), these latter defects in *pmr1*Δ cells were largely suppressed by loss of Smf2. Thus, our findings posit a model in which elevated cytoplasmic Mn^2+^ levels antagonize autophagy, mitophagy, and Rtg1-3-dependent retrograde signaling presumably through activation of TORC1.

### *MnCl_2_ stimulates TORC1 kinase activity* in vivo *and* in vitro

The yeast Tor1 kinase is a member of the phosphatidylinositol 3-kinase (PI3K) -related kinase (PIKK) family that can phosphorylate the human eukaryotic translation initiation factor 4E binding protein (eIF4-BP/PHAS-I) *in vitro* (43). Curiously, it does so much more efficiently when the respective *in vitro* kinase assays contain Mn^2+^ rather than Mg^2+^ as the sole divalent cation, a property that it appears to share with PI3-kinases (44–46). A similar preference for Mn^2+^ over Mg^2+^ has also been observed in mTOR kinase autophosphorylation assays (31, 32). Based on these and our observations, we decided to assess whether Mn^2+^ may act as a metal cofactor for TORC1 activity *in vitro* using TORC1 purified from yeast and a truncated form of Lst4 (Lst4^Loop^; (47)) as a substrate. In control experiments without divalent ions, TORC1 activity remained undetectable (Figure 3A and B). The addition of MnCl_2_, however, not only stimulated TORC1 *in vitro* in a concentration-dependent manner but also activated TORC1 dramatically more efficiently than MgCl_2_ (with 25-fold lower levels of MnCl_2_ [38 μM] than MgCl_2_ [980 μM] promoting half-maximal activation of TORC1; Figure 3A and B). We next considered the possibility that Mn^2+^ is superior to Mg^2+^ in favoring the coordination of ATP in the catalytic cleft or TORC1. Supporting this idea, we found that Mn^2+^ significantly reduced the K_m_ for ATP (5.3-fold) of TORC1 (Figure 3C and 3D). Moreover, even in the presence of saturating Mg^2+^ levels (*i.e*. 4 mM), the addition of 160 μM Mn^2+^ was able to enhance the V_max_ almost 2-fold and decrease the K_m_ for ATP of TORC1 from 50.7 to 34.4 μM, which indicates that Mn^2+^ can efficiently compete with Mg^2+^ and thereby activate TORC1. Notably, the conditions used in our *in vitro* kinase assays are quite comparable to the *in vivo* situation: accordingly, intracellular Mg^2+^ levels in yeast are approximately around 2 mM (48), while the Mn^2+^ levels range from 26 μM in WT cells to 170 μM in *pmr1*Δ cells (49). Our *in vitro* assays, therefore, provide a simple rationale for why *pmr1*Δ cells are resistant to rapamycin: elevated Mn^2+^ levels in *prm1*Δ favorably boost the kinetic parameters of TORC1.

**Figure 3.**
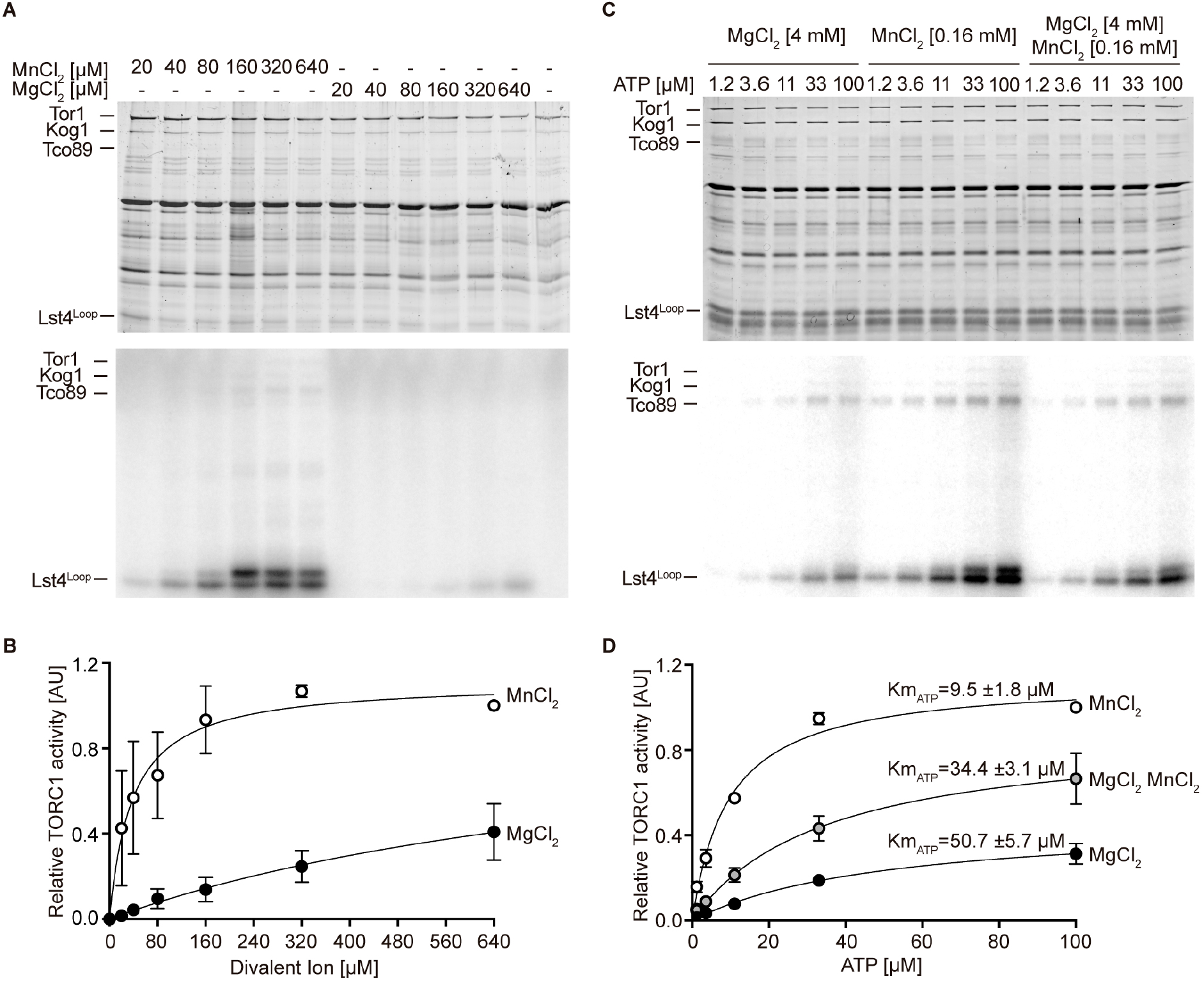
MnCl_2_ stimulates TORC1 kinase activity *in vitro* and *in vivo*. (**A**) *In vitro* TORC1 kinase assays using [γ-^32^P]-ATP, recombinant Lst4^Loop^ as substrate, and increasing concentrations (2-fold dilutions) of MgCl_2_ or MnCl_2_. Substrate phosphorylation was detected by autoradiography (lower blot) and Sypro Ruby staining is shown as loading control (upper blot). (**B**) Quantification of the assay shown in (A). Curve fitting and parameter calculations were performed with GraphPad Prism. Data shown are means (± SEM, n = 3). (**C**) *In vitro* kinase assays (as in A) using the indicated concentrations of MgCl_2_ and/or MnCl_2_ and increasing concentrations of ATP. Substrate phosphorylation was detected by autoradiography (lower blot) and Sypro Ruby staining is shown as loading control (upper blot). (**D**) Quantification of the assay shown in (C). Curve fitting and parameter calculations were performed with GraphPad Prism. Data shown are means (± SEM, n = 3).

### TORC1 regulates NRAMP transporter protein levels

To maintain appropriate intracellular Mn^2+^ concentrations, cells adjust Smf1 and Smf2 protein levels through Bsd2-dependent and -independent, post-translational modifications that ultimately trigger their vacuolar degradation through the endocytic multivesicular body pathway (12, 50). Because NRAMP transporters are important for Mn-dependent TORC1 activation, and because TORC1 is often embedded in regulatory feedback loops to ensure cellular homeostasis (51), we next asked whether rapamycin-mediated TORC1 inactivation may affect the levels and/or localization of Smf1 and Smf2 using strains that express GFP-Smf1 or Smf2-GFP. Rapamycin treatment triggered a strong increase in GFP-Smf1 levels (Figure 4A and 4B), which is likely due to transcriptional activation of Smf1 under these conditions as published earlier (52). In line with this interpretation, we found the respective increase to be abolished when rapamycin-treated cells were co-treated with the protein synthesis inhibitor cycloheximide (CHX). In addition, because GFP-Smf1 appeared to be degraded at a similar rate in CHX-treated cells that were treated, or not, with rapamycin, our data further indicate that TORC1 does not control Smf1 levels through posttranslational control of Smf1 turnover. Interestingly, GFP-Smf1 remained predominantly at the plasma membrane in rapamycin-treated cells, even though these cells also accumulated more cleaved GFP in the vacuoles (Figure 4C). We infer from these data that TORC1 inhibition activates Smf1 expression, likely as part of a feedback control loop through which TORC1 couples its activity to Mn^2+^ uptake.

**Figure 4.**
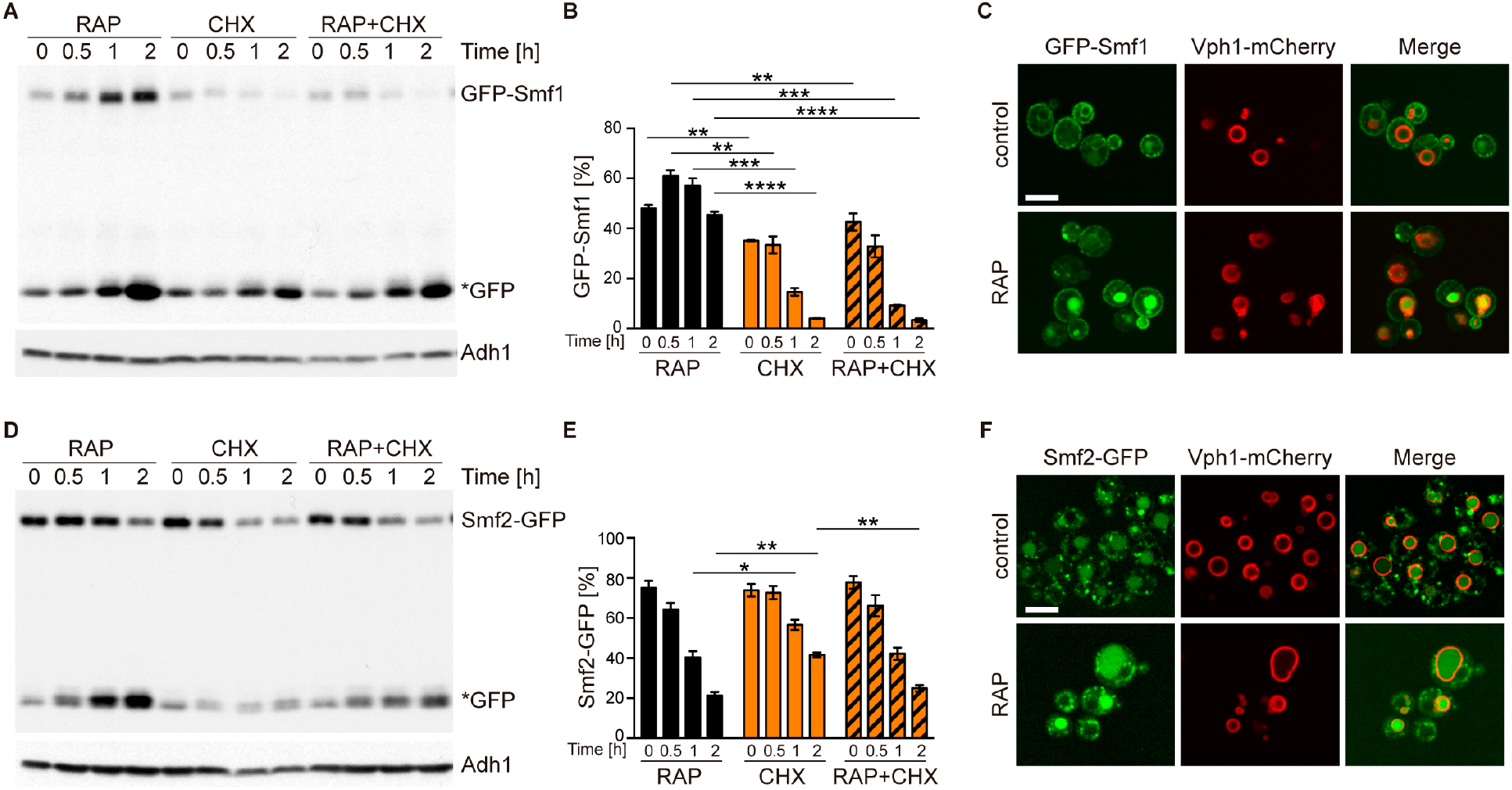
TORC1 regulates NRAMP transporter levels. (**A**) GFP-Smf1 expressing cells were cultivated for 5-6 h in a synthetic medium devoid of manganese sulfate before being treated with 200 ng/ml rapamycin (RAP), 25 μg/ml cycloheximide (CHX) or both compounds (RAP+CHX) for the indicated times. GFP-Smf1 and cleaved GFP (*GFP) protein levels were analyzed by immunoblotting using an anti-GFP antibody. Alcohol dehydrogenase (Adh1) protein levels, probed with anti-Adh1 antibodies, served as a loading control. (**B**) Quantification of GFP-Smf1. Percentages of GFP-Smf1 relative to the total GFP signal (GFP-Smf1 + GFP). Data represent means ± SEM of independent experiments (*n*=3). Statistical analysis: unpaired two-tailed t-test. * p<0.05; ** p<0.01, *** p<0.001, **** p<0.0001. **(C)**Microscopic analysis of GFP-Smf1 localization. Cells co-expressing GFP-Smf1 and the vacuolar marker Vph1-mCherry were grown exponentially in a manganese-free medium for 5-6 h, then treated with 200 ng/ml rapamycin for 2 h. Scale bar represents 5 μm. (**D-E**) Cells expressing Smf2-GFP from its endogenous locus were grown, treated, and processed as in (A). (**F**) Microscopic analysis of Smf2-GFP localization. Cells co-expressing Smf2-GFP and the vacuolar marker Vph1-mCherry were cultivated, treated, and examined as in (C).

Rapamycin-treated cells that were either co-treated or not with CHX exhibited Smf2-GFP levels that steadily decreased at a similar rate, with cleaved GFP accumulating in parallel (Figure 4D and 4E). Since this rate appeared to be higher than the one observed in cells treated with CHX alone, our data suggest that TORC1 antagonizes the turnover of Smf2. This was further corroborated by our fluorescence microscopy analyses revealing that the GFP signal in Smf2-GFP-expressing cells shifted from late Golgi/endosomal foci (see (37)) to a predominant signal within the vacuolar lumen when cells were treated with rapamycin (Figure 4F). Whether this event is an indirect consequence of higher Mn^2+^ uptake under these conditions (see above), or potentially part of a local feedback control mechanism of endosomal TORC1 (53), remains to be addressed in future studies.

### Mn^2+^ activates mTORC1 signaling in human cells

To assess whether the response of TORC1 to Mn^2+^ is conserved among eukaryotes, we investigated if externally supplied MnCl_2_ activates mTORC1 signaling in mammalian cells. For this purpose, we used the human cell lines U2OS and HEK293T, which are widely used for the cellular dissection of mTORC1-mediated mechanisms in response to nutritional inputs (54). First, we examined the sufficiency of Mn^2+^ to maintain the activity of the mTORC1 pathway by incubating U2OS cells in an amino acid starvation medium supplemented with increasing amounts of MnCl_2_ (0-1 mM) for 2h. mTORC1 activity was assessed by monitoring phosphorylation of the mTORC1-downstream targets RPS6KB (ribosomal protein S6 kinase B; phospho-Threonine^389^) and RPS6 (ribosomal protein S6; phospho-Serine^235-236^). As shown in Figure 5A, and as expected, both RPS6KB and RPS6 were fully phosphorylated at those specific residues in cells incubated in the presence of amino acids and completely de-phosphorylated in cells incubated in the absence of amino acids, without MnCl_2_. In agreement with a positive action of Mn^2+^ towards mTORC1 in human cells, we observed a robust and dose-dependent increase in both RPS6KB and RPS6 phosphorylation in amino acid-starved cells incubated in the presence of MnCl_2_ at concentrations higher than 0.05 mM, with maximum phosphorylation observed at 0.5 mM of MnCl_2_. Confirming this result, we observed a similar response of RPS6KB and RPS6 phosphorylation to MnCl_2_ treatment in HEK293T cells, except that the maximum response was reached at 1 mM (Figure 5B). These results indicate that Mn^2+^ is sufficient to maintain the activity of the mTORC1 pathway in the absence of amino acid inputs, at least during short periods (2 h). To corroborate this conclusion, we also tested the phosphorylation of additional downstream targets of mTORC1 in response to a fixed concentration of MnCl_2_ (0.35 mM) in U2OS cells. Amino acid-starvation of U2OS cells with MnCl_2_ was sufficient to maintain the phosphorylation of the downstream targets of mTORC1 RPS6KB, RPS6, EIF4EBP1, and ULK1 (unc-51 like autophagy activating kinase 1) (Figure 5C). In line with our finding that MnCl_2_ retained AKT1 (AKT serine/threonine kinase 1) phosphorylation in amino-acid starved cells, Mn^2+^ has been reported to activate the mTORC1 upstream kinase AKT1, which is also a known mTORC2 downstream target (55, 56). However, AKT1 phosphorylation appeared to be much less pronounced than the phosphorylation of the mTORC1 downstream targets, validating the specificity of Mn towards mTORC1. Whether Mn^2+^ may stimulate TORC2 activity remains to be explored.

**Figure 5.**
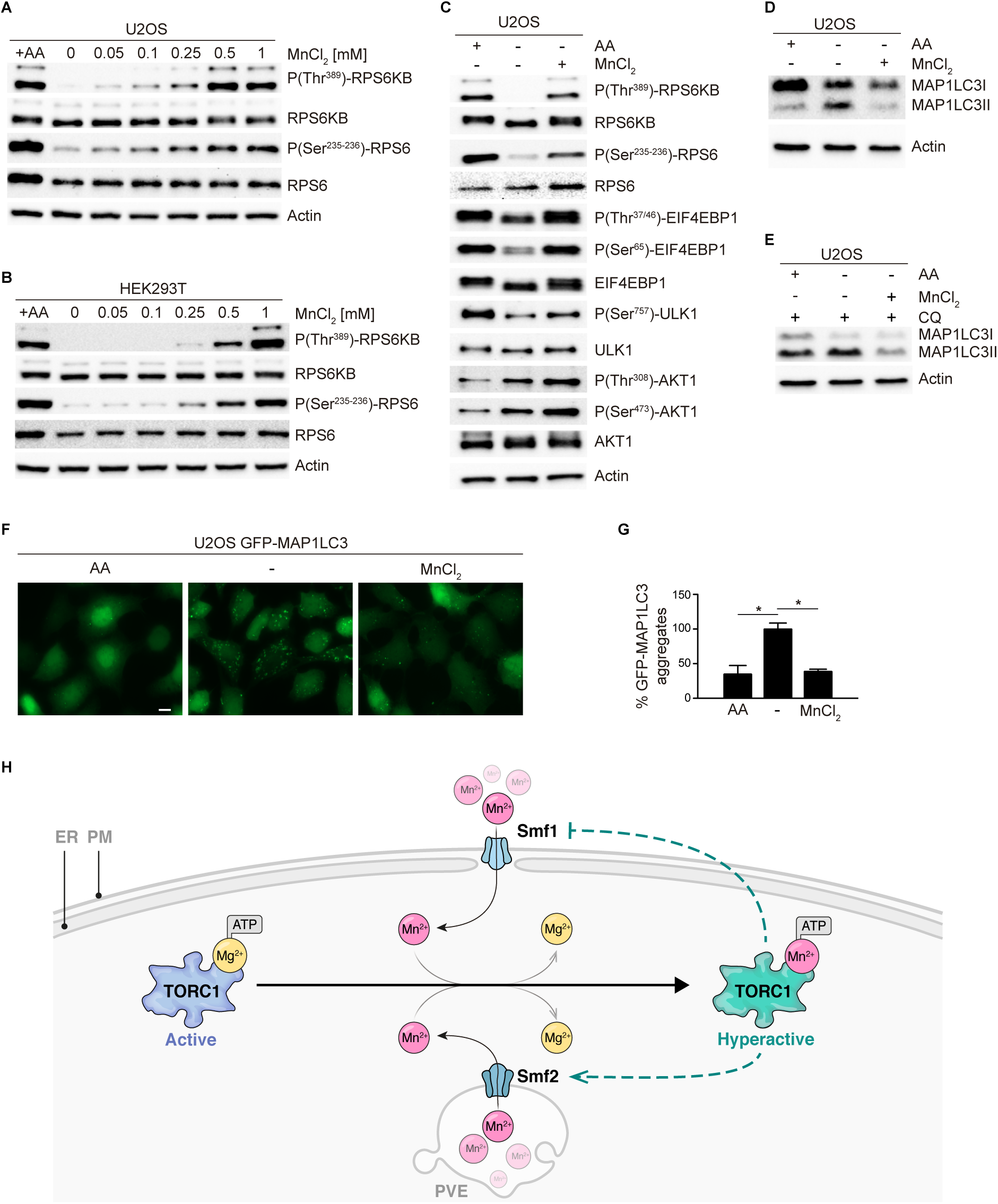
MnCl_2_ maintains mTORC1 activity during starvation in human cells. (**A-B**) Human U2OS (A) and HEK293T (B) cells were starved for all amino acids and supplemented with increasing concentrations of MnCl_2_ as indicated for 2 hours. Phosphorylation of the mTORC1 downstream targets RPS6KB and RPS6 was assessed by immunoblot analysis. A control with cells incubated in the presence of all proteinogenic amino acids (+AA) was included as a positive control. (**C**) Human U2OS were starved for all amino acids (-AA) and supplemented with 0.35 mM of MnCl_2_ for 2 h. Phosphorylation of RPS6KB, RPS6, EIF4EBP1, ULK1, and AKT1 was assessed by immunoblot analysis. A control with cells incubated in the presence of all proteinogenic amino acids (+AA) was included as a positive control. (**D-E**) Human U2OS cells were treated as in (C) for 4 h either in the absence (D) or the presence (E) of chloroquine (CQ). Autophagic marker MAP1LC3I/II was then analyzed by immunoblot. (**F-G**) GFP-MAP1LC3 expressing U2OS cells were incubated as in (D). GFP-MAP1LC3 aggregation was assessed by confocal microscopy (F) and quantified using ImageJ software (G).Scale bar indicates 10 μm. Values were normalized to -AA condition. Graphs represent mean ± S.D., with n=3 (*, p<0.05, ANOVA followed by Tukey’s test). (**H**) Model of Mn^2+^-driven TORC1 activation. The Smf1/2 NRAMP transporter-dependent increase in cytoplasmatic Mn^2+^ levels favors TORC1-Mn^2+^ binding and ATP coordination, leading to TORC1 hyperactivation. NRAMP transporters are part of a feedback control mechanism impinged by TPRC1 (dashed lines).

Finally, to assess the physiological relevance of mTORC1 activation in response to Mn^2+^, we analyzed the effect of MnCl_2_ treatment in autophagy, a cellular process inhibited by TORC1 both in yeast and mammalian cells (57, 58). To this end, we analyzed the autophagic marker MAP1LC3I/II (microtubule-associated protein 1 light chain 3) (59). During autophagy initiation, MAP1LC3I is lipidated, thereby increasing the levels of MAP1LC3II. As previously reported, the withdrawal of amino acids led to a rapid increase in MAP1LC3II levels, indicating an increase in autophagy initiation (Figure 5D). Of note, the addition of MnCl_2_ completely abolished the increase in MAP1LC3II levels, thus confirming that Mn^2+^prevented the initiation of autophagy downstream of mTORC1. These results were also observed in cells incubated in the presence of chloroquine, an inhibitor of the fusion of autophagosomes with the lysosome, thus confirming that Mn^2+^ influences autophagy initiation, the process controlled by mTORC1 (Figure 5E). In addition, we analyzed the aggregation of MAP1LC3 upon autophagosome formation in U2OS cells stably expressing a GFP-MAP1LC3 construct. GFP aggregation indicates autophagosome formation in these cells. Similar to what we observed with endogenous MAP1LC3, GFP aggregation was increased during amino acid withdrawal and this increase was completely abolished by MnCl_2_ treatment (Figure 5F-G). This result further confirms that the Mn-dependent activation of mTORC1 in human cells is physiologically relevant for autophagy inhibition. Altogether, our results show that Mn^2+^ is sufficient to activate mTORC1 in the absence of other inputs in human cells and to inhibit autophagy downstream of mTORC1. These findings indicate that Mn^2+^-mediated activation of TORC1 is evolutionarily conserved from yeast to humans.

## Discussion

Studies with yeast lacking the divalent Ca/Mn ion Golgi transporter Pmr1, which displays an increase in intracellular Mn^2+^ levels due to impaired Mn^2+^ detoxification (49), have pinpointed a functional link between Mn^2+^ homeostasis and TORC1 signaling (33, 34). However, the underlying mechanism(s) of how Mn^2+^impinges on TORC1 has so far remained elusive. Our study identifies Mn^2+^ as a divalent metal cofactor that stimulates the enzymatic activity of the TORC1 complex *in vitro* and suggests that Mn^2+^ functions similarly *in vivo* both in yeast and mammalian cells. Our results support a model in which the PI3K-related protein kinase Tor1 is activated by Mg^2+^, but requires Mn^2+^ as a metal cofactor for maximal activity as this allows it to better coordinate ATP (Figure 5H). While it seemingly shares this feature with PI3Ks (44–46), serine/threonine protein kinases are generally known to be preferably activated by Mg^2+^ rather than Mn^2+^ (60). We deem it therefore possible that the respective Mn^2+^-driven activation of TORC1 is more specifically related to the architecture of its catalytic center. Resolution of this issue will therefore require future molecular dynamic modeling and/or structural analyses.

Our current study also highlights a need for coordinated control of TORC1 activity and cytoplasmic Mn^2+^ levels. The latter critically depend on a set of conserved NRAMP transporters, such as yeast Pmr1 and Smf1/2, and their mammalian homologs ATP2C1/SPCA1 (38) and SLC11A2 (DMT1; (61) and this work). In yeast, cytoplasmic Mn levels can be increased by the loss of Pmr1, which causes a defect in sequestration of Mn^2+^ in the Golgi. Alternatively, they can also be boosted by increased Smf1-mediated Mn^2+^ uptake combined with Smf2-mediated Mn^2+^ depletion in endosomes following the loss of Bsd2, an adaptor protein for the NEDD4 family E3 ubiquitin ligase Rsp5 (12) that normally favors vacuolar degradation of Smf1/2 (62, 63). In both cases, *i.e.* loss of Pmr1 or of Bsd2 (specifically upon addition of Mn^2+^ in the medium), enhanced cytoplasmic Mn^2+^ levels mediate higher TORC1 activity and hence the resistance of cells to low doses of rapamycin. Conversely, our expression and localization studies of GFP-Smf1 and Smf2-GFP indicate that TORC1, while being controlled by Mn^2+^ levels, also employs feedback control mechanisms that regulate Smf1 expression and Smf2 turnover. These observations are consistent with the recent paradigm shift assigning TORC1 an essential role in cellular and organismal homeostasis (51) and extend this role of TORC1 to Mn metabolism. A more detailed understanding of these regulatory circuits involving TORC1 and NRAMP transporters (and *vice versa*) will, however, require future analyses of the specific subcellular pools of Mn^2+^ within different organelles using electron-nuclear double resonance spectroscopy (49) or inductively coupled plasma mass spectrometry (64). Of note, Mn^2+^ also acts as a cofactor for glutamine synthetase (3) that produces the TORC1-stimulating amino acid glutamine (65). Hence, Mn^2+^ homeostasis, amino acid metabolism, and TORC1 may be subjected to even more intricate, multilayered feedback regulatory circuits.

Our finding that Mn^2+^ activates TORC1/mTORC1 may have important implications in different biological contexts. For instance, previous work has shown that chronic exposure to high levels of Mn^2+^causes autophagic dysfunction and hence accumulation of compromised mitochondria in mammalian astrocytes due to reduced nuclear localization of TFEB (transcription factor EB), a key transcription factor that coordinates the expression of genes involved in autophagy (66). Because mTORC1 inhibits TFEB (67) and mitochondrial quality can be improved by rapamycin-induced TFEB induction (and consequent stimulation of autophagy) (68), our study now provides a simple rationale for the observed accumulation of damaged mitochondria upon Mn^2+^ exposure. Accordingly, excess Mn^2+^ antagonizes autophagy and mitophagy at least in part through mTORC1 activation and subsequent TFEB inhibition, which prevents proper disposal of hazardous and reactive oxygen species-producing mitochondria. This model also matches well with our previous observation that rapamycin restricts cell death associated with anomalous mTORC1 hyperactivation (58). Finally, our data indicate that Mn^2+^-driven TORC1 activation and the ensuing inhibition of auto- and mitophagy are also employed by yeast cells, which highlights the primordial nature of these processes.

Another area where our findings may be relevant relates to the use of Mn^2+^ to stimulate antitumor immune responses. Recent studies have shown that Mn^2+^ is indispensable for the host defense against cytosolic, viral double-stranded DNA as it mediates activation of the DNA sensor CGAS (cyclic GMP-AMP synthase) and its downstream adaptor protein STING1 (stimulator of interferon response cGAMP interactor 1) (69). Since Mn^2+^ stimulates the innate immune sensing of tumors, Mn administration has been suggested to provide an anti-tumoral effect and improve the treatment of cancer patients (70). Nevertheless, mTOR (mechanistic target of rapamycin kinase) hyperactivation is known to promote tumor progression (71), and carcinoma and melanoma formation have previously been associated with mutations in the human *PMR1* ortholog *ATP2C1* that cause Hailey-Hailey disease (72). Based on our findings, it is therefore possible that tumor formation in these patients may be causally linked to Mn^2+^-dependent DNA replication defects (16), stress response (73), and mTORC1 activation.

In addition to the potential relevance of our findings in the context of chronic Mn^2+^ exposure and immune stimulation, our findings may also provide new perspectives to our understanding of specific neuropathies. For instance, exposure to Mn dust or Mn containing smoke, as a byproduct of metal welding, is well known to cause a parkinsonian-like syndrome named manganism, a toxic condition of Mn poisoning with dyskinesia (74). Interestingly, while dyskinesia has been connected to L-dopamine-mediated activation of mTORC1 (75), our findings suggest that Mn^2+^-driven mTORC1 hyperactivation may impair autophagy and thereby contribute to neurological diseases (76). In line with this reasoning, compounds that inhibit TORC1 activity, and thus stimulate autophagy, have been suggested as therapeutics for the treatment of Parkinson-like neurological symptoms (77). In this context, Huntington disease (HD) is another example of patients suffering from a proteinopathy characterized by parkinsonian-like neurological syndromes (78). In HD patients, expansion of the polyglutamine tract in the N-terminus of the huntingtin protein leads to protein aggregation (79) and, intriguingly, HD patients exhibit reduced Mn^2+^ levels in the brain (80). This begs the question of whether the respective cells aim to evade Mn-driven mTORC1 activation, *e.g*. by reducing Mn uptake or sequestration of Mn^2+^ within organelles, to stimulate huntingtin protein degradation via autophagy. Finally, Mn^2+^ also contributes to prion formation in yeast (81), and elevated Mn^2+^ levels have been detected in the blood and the central nervous system of Creutzfeldt-Jakob patients (82). It will therefore be exciting to study the cell type-specific impact of Mn^2+^-driven mTORC1 activation on metabolism, genome stability, checkpoint signaling, and the immune response, all processes that play a key role in neurological diseases and aging-related processes.

## Materials and Methods

### Antibodies

Primary antibodies against GFP (Clontech, JL-8; 1:5000), G-6-PDH (Sigma-Aldrich, A9521; 1:5000), Adh1 (Calbiochem, 126745; 1:200000), RPS6 (Cell Signalling Technology, 2217; 1:1000), phospho-RPS6 (Ser^235-236^) (Cell Signalling Technology, 4856; 1:1000), RPS6KB (Cell Signalling Technology, 2708; 1:1000), phospho-RPS6KB (Thr^389^) (Cell Signalling Technology, 9205; 1:1000), EIF4EBP1 (Cell Signalling Technology, 9452; 1:1000), phospho-EIF4EBP1 (Thr^37/46^) (Cell Signalling Technology, 2855; 1:1000), phospho-EIF4EBP1 (Ser^65^) (Cell Signalling Technology, 9451; 1:1000), AKT1 (Cell Signalling Technology, 4691; 1:1000), phospho-AKT(Ser^473^) (Cell Signalling Technology, 4060; 1:1000), phospho-AKT(Thr^308^) (Cell Signalling Technology, 13038; 1:1000), ULK1 (Cell Signalling Technology, 8359; 1:1000), phospho-ULK1 (Cell Signalling Technology, 8359; 1:1000), MAP1LC3 AB (Cell Signalling Technology, 12741; 1:1000), β-actin (Cell Signalling Technology, 4967; 1:1000) were used.

### Yeast strains, Plasmids, and Growth Conditions

Yeast strains and plasmids used in this study are listed in Tables S2 and S3. Gene disruption and tagging were performed with standard high-efficiency recombination methods. To generate strain MP6988, pSIVu-SMF1p-GFP-SMF1 was digested with *Pac*I and transformed into *smf1*Δ strain YOL182C. Yeast cells were grown to mid-log phase in synthetic defined medium (SD; 0.17% yeast nitrogen base, 0.5% ammonium sulfate, 2% glucose, 0.2% drop-out mix) at 30°C. To induce autophagy, mitophagy, and Rtg1-Rtg3 retrograde signaling, cells were treated with 200 ng/ml rapamycin (Biosynth Carbosynth, AE27685) for the indicated time. For telomere length analyses, cells were grown for 3 days in rich medium (YPAD; 1% yeast extract, 2% peptone, 2% glucose, 0.004% adenine sulfate) with or without the addition of 10 mM CaCl_2_. For imaging of Rtg3-GFP, cells were grown at 26°C in SD-MSG-ura medium (0.17% yeast nitrogen base, 0.5% monosodium glutamate, 2% glucose, 0.2% drop-out mix -ura) to exponential phase and treated or not with 200 ng/ml rapamycin for 30 min. For GFP1-Smf1 and Smf2-GFP analyses, cells were grown in SD-ura medium. Overnight precultures were quickly spun and diluted into SD-ura medium devoid of manganese sulfate (0.19% yeast nitrogen base without manganese sulfate (Formedium; CYN2001), 0.5% ammonium sulfate, 2% glucose, and 0.2% drop-out mix -ura). After 5 to 6 h of growth, cells were treated with either rapamycin (200 ng/ml), cycloheximide (25 μg/ml), or both compounds for the indicated times.

### Yeast Cell Lysate Preparation and Immunoblotting

Cells grown to mid-log phase were treated with 6.7% trichloroacetic acid (TCA, final concentration), pelleted, washed with 99% acetone, dried, dissolved in urea buffer (6 M urea, 50 mM Tris-HCl pH 7.5, 1% SDS, 1 mM PMSF and 10 mM NaF) and disrupted with glass beads using a Yasui Kikai homogenizer. Cell lysates were heated at 65° C for 10 min in Laemmli SDS sample buffer, centrifugated at 15’000 g for 1 min, and the supernatants were subjected to SDS-PAGE and immunoblotted. Heat denaturation of samples was omitted to detect GFP-Smf1 and Smf2-GFP. Chemiluminescence signals were captured in an Amersham ImageQuant 800 Imager and quantified with ImageQuant TL software (Cytiva).

### Drug Sensitivity Assays

Yeast cells grown to mid-log phase were adjusted to an initial A600 of 0.2, serially diluted 1:10, and spotted onto plates without or with rapamycin at the indicated concentrations (see figure legends). 1 mM MnCl_2_ and/or 10 mM CaCl_2_ were added to the medium when indicated. Plates were then incubated at 30°C for 3 to 4 days.

### Analysis of Telomere Length

Genomic DNA was isolated from yeast strains grown in YPAD for 3 days with or without the addition of 10 mM CaCl_2_. DNA was digested with *Xho*I, separated on a 1% agarose-Tris borate EDTA gel, transferred to a Hybond XL (Amersham Biosciences) membrane, and hybridized with a ^32^P-labeled DNA probe specific for the terminal Y’ telomere fragment. The probe was generated by random hexanucleotide-primed DNA synthesis using a short Y’ specific DNA template, which was generated by PCR from genomic yeast DNA using the primers Y up (5’-TGCCGTGCAACAAACACTAAATCAA-3’) and Y’ low (5’-CGCTCGAGAAAGTTGGAGTTTTTCA-3’). Three independent colonies of each strain were analyzed to ensure reproducibility.

### *In vitro* TORC1 kinase assays

TORC1 was purified and kinase assays were performed as previously described (47) with minor modifications. For the kinase assays in the presence of various concentrations of magnesium or manganese, reactions were performed in a total volume of 20 μl with kinase buffer (50 mM HEPES/NaOH [pH 7.5], 150 mM NaCl), 400 ng of purified His_6_-Lst4^Loop^, 60 ng TORC1 (quantified based on the Tor1 subunit) and 640, 320, 160, 80, 40 or 20 μM MgCl_2_/MnCl_2_. The reaction was started by adding 1.4 μl of ATP mix (18 mM ATP, 3.3 mCi [γ-^32^P]-ATP [Hartmann Analytic, SRP-501]). For the kinase assays in the presence of various concentrations of ATP, reactions containing 4 mM MgCl_2_, 160 μM MnCl_2_, or both were started by adding 1.4 μl of stock ATP mix (5.7 mM ATP, 3.3 mCi [γ-^32^P]-ATP [Hartmann Analytic, SRP-501]) or two-fold serial dilutions. After the addition of sample buffer, proteins were separated by SDS-PAGE, stained with SYPRO Ruby (Sigma-Aldrich, S492) (loading control), and analyzed using a phosphoimager (Typhoon FLA 9500; GE Healthcare). Band intensities were quantified with ImageJ and data were analyzed with GraphPad Prism using the Michaelis-Menten non-linear fitting.

### Cell culture

U2OS and HEK293T cell lines were obtained from the American Type Culture Collection (ATCC). GFP-MAP1LC3 expressing U2OS cells were kindly provided by Dr. Eyal Gottlieb (Cancer research UK, Glasgow, UK). All the cell lines were grown in high glucose DMEM base medium (Sigma-Aldrich, D6546) supplemented with 10 % of fetal bovine serum, glutamine (2 mM), penicillin (Sigma-Aldrich, 100 μg/ml), and streptomycin (100 μg/ml), at 37 °C, 5 % CO_2_ in a humidified atmosphere. Amino acid starvation was performed with EBSS medium (Sigma-Aldrich, E2888) supplemented with glucose at a final concentration of 4.5 g/L. When indicated, the starvation medium was complemented with MnCl_2_ to a final concentration of 0.05 −1 mM, chloroquine (Sigma-Aldrich; 10 μM), and amino acids, by adding MEM Amino Acids (Sigma-Aldrich, M5550), plus MEM Non-essential Amino Acid Solution (Sigma-Aldrich, M7145) and glutamine (2 mM).

### Confocal and fluorescence microscopy

Yeast cells: For imaging of GFP-Tor1, GFP-Smf1 and Smf2-GFP, images were captured with an inverted spinning disk confocal microscope (Nikon Ti-E, VisiScope CSU-W1, Puchheim, Germany) that was equipped with an Evolve 512 EM-CDD camera (Photometrics), and a 100x 1.3 NA oil immersion Nikon CFI series objective (Egg, Switzerland). Images were then processed using the FIJI-ImageJ software. For imaging of Rtg3-GFP, images were obtained by projection of a focal plane image derived from wide-field fluorescence microscopy (DM-6000B, Leica) at 100x magnification using L5 filters and a digital charge-coupled device camera (DFC350, Leica). Pictures were processed with LAS AF (Leica).

Human cells: 8 x 10^5^ cells were grown in coverslips for 24h and treated for 4h as indicated. Thereafter, cells were fixed with 4% paraformaldehyde (Sigma-Aldrich) in PBS for 10 min at room temperature. For GFP-MAP1LC3 assessment, after three washes with PBS, coverslips were mounted with Prolong containing DAPI (Invitrogen). Samples were imaged with a Zeiss Apotome microscope. GFP aggregation in microscopy images was assessed using ImageJ software.

### Statistical analyses

Statistical analyses were performed using GraphPad Prism 7.0. Statistical significance was determined from at least 3 independent biological replicates using either Student’s t-test, or ANOVA followed by Tukey’s multiple comparison test. Differences with a P-value lower than 0.05 were considered significant. * p<0.05; ** p<0.01; *** p<0.001; **** p<0.0001. The number of independent experiments (n), specific statistical tests and significance are described in the figure legends.

## Acknowledgments

We thank María Díaz de la Loza for scientific illustration work, and V. Albanèse, E. de Nadal, V. Goder, K. D. Hirschi, C. Ungermann, and E. Gottlieb for plasmids, yeast strains, and cell lines. Research was funded by grants from the University of Seville (2020/00001326), Junta de Andalucía/European Union Regional Funds (P20-RT-01220) and EMBO (STF-8685) to R.E.W.; the Swiss National Science Foundation (310030_166474/184671) to C.D.V.; the Spanish Ministry of Science, Innovation and Universities (PGC2018-096244-B-I00) to R.D. The author L.Z. was the recipient of a predoctoral grant from the Spanish Ministry of Science, Innovation and Universities (FPU19/04914).

