## supplementary for "Manganese is a Physiologically Relevant TORC1 Activator in Yeast and Mammals"

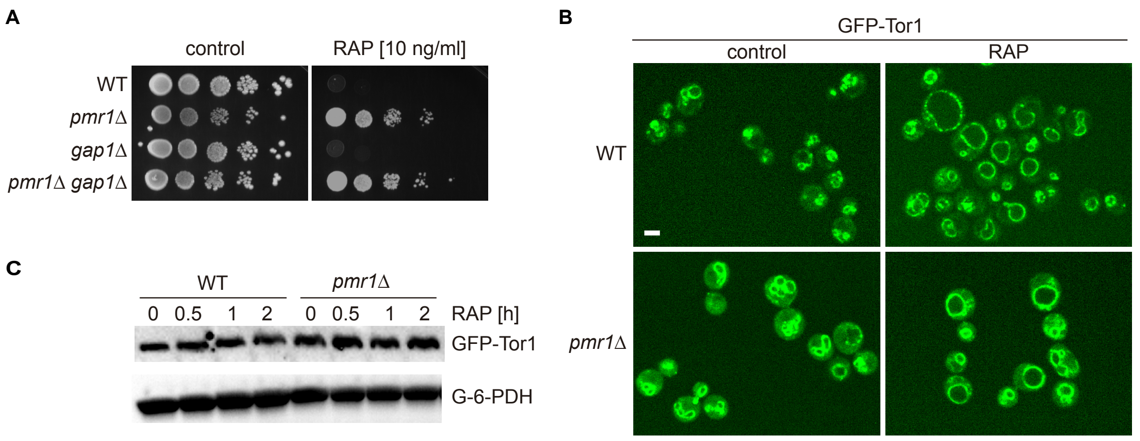


**Figure S1.** *pmr1Δ* rapamycin resistance is not linked to Gap1 or Tor1 localization. (**A**) Sensitivity of indicated strains to rapamycin (RAP). 10-fold dilutions of exponentially growing cells are shown. (**B**) Spinning disk microscopic analysis of GFP-Tor1 localization WT and *pmr1Δ* cells upon treatment with 200 ng/ml rapamycin (+RAP) for 30 minutes. Scale bar represents 5 µm. (**C**) Exponentially growing cells expressing GFP-Tor1 from the endogenous locus were treated for up to 2 h with 200 ng/ml rapamycin (RAP). GFP-Tor1 protein levels were analyzed by immunoblotting with an anti-GFP antibody. G-6-PDH levels were used as a loading control.

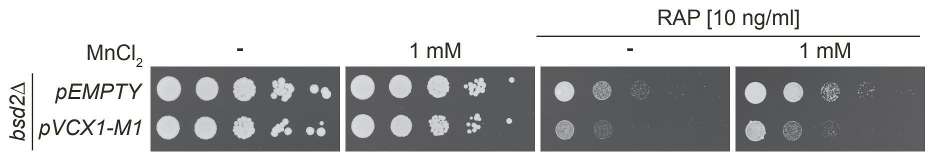


**Figure S2:** Growth of *bsd2Δ* mutants expressing Vcx1-M1 from plasmid *pVCX1-M1* on $MnCl_2$ and/or rapamycin (RAP) containing medium. 10-fold dilutions of exponentially growing cells are shown. Compound concentrations are indicated.

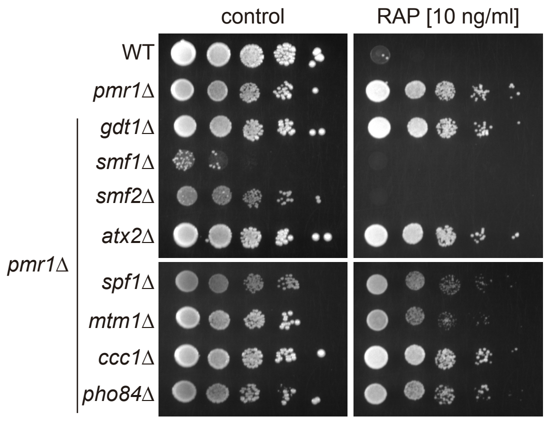


**Figure S3:** NRAMP transporters mediate rapamycin resistance of *pmr1*Δ mutants. Sensitivity of the indicated strains to rapamycin (RAP). 10-fold dilution of exponentially growing cells are shown. Data obtained in medium supplemented with $CaCl_2$ is shown in Figure 1D. Note that growth of the *pmr1*Δ *smf1*Δ double mutant is strongly impaired without $CaCl_2$ addition.

**Table S1.** RTG1-3 target genes down-regulated in *pmr1*Δ cells.

| Gene name | Fold change |
|---|---|
| *BDH2* | -4,8 |
| *CIT2* | -1,65 |
| *FMP48* | -5,1 |
| *HSP12* | -4,75 |
| *HXT5* | -2,2 |
| *IDH1* | -1,85 |
| *IDH2* | -1,6 |
| *MSC1* | -2,75 |
| *PHM7* | -1,85 |
| *RTC3* | -2,2 |
| *SPS100* | -3,15 |
| *TKL2* | -3 |
| *YNL194C* | -8,4 |

Data from García-Rodríguez et al, 2012. GEO accession GSE29420.

**Table S2.** Plasmids used in this study.

| Plasmid | Relevant Genotype | Source |
|---|---|---|
| VGp160 | *GFP-ATG8 CEN URA3* | V. Goder |
| pSIVu | Cloning vector for single integration into the *URA3* locus | S. Pelet |
| pVA1458 | *SMF1pGFP-SMF1 URA3* | V. Albanese/S. Leon |
| p4301 | *SMF1pGFP-SMF1 Sac*I/*Kpn*I fragment from pVA1458 cloned into single integration plasmid pSIVu *URA3* | This study |
| p2GPD-SMF2 | *SMF2 URA3* | This study |
| p2GPD-Slc11a1 | *cSlc11a1 URA3* | This study |
| p2GPD-Slc11a2 | *cSlc11a2 URA3* | This study |
| pRS416-RTG3 | *RTG3p::RTG3-GFP URA3* | E. de Nadal |
| pRS416 | *URA3* | P. Hieter |
| p2GPD | *URA3* | K. D. Hirschi |
| pVCX1-M1 | *VXC1-M1 URA3* | K. D. Hirschi |

**Table S3.** Yeast strains used in this study.

| Strains | Relevant Genotype | Source |
|---|---|---|
| BY4741 | *MATa his3Δ1 leu2Δ0 met15Δ0 ura3Δ0* | Euroscarf |
| YGL167C | *pmr1Δ::KAN*, isogenic to BY4741 | Euroscarf |
| YBR290W | *bsd2Δ::KAN*, isogenic to BY4741 | Euroscarf |
| RW187 | *pmr1Δ::NAT bsd2Δ::KAN*, isogenic to BY4741 | This study |
| YOL122C | *smf1Δ::KAN*, isogenic to BY4741 | Euroscarf |
| NG222 | *pmr1Δ::NAT smf1Δ::KAN*, isogenic to BY4741 | R. Wellinger |
| YHR050W | *smf2Δ::KAN*, isogenic to BY4741 | Euroscarf |
| NGY183 | *pmr1Δ::NAT smf2Δ::KAN*, isogenic to BY4741 | R. Wellinger |
| RWY188 | *bsd2Δ::KAN smf2Δ::NAT*, isogenic to BY4741 | This study |
| RKH395 | *MATa LEU2::GFP-TOR1 his3Δ1 leu2Δ0 ura3Δ0* | C. De Virgilio |
| RWY128 | *pmr1Δ::NAT,* isogenic to RKH395 | This study |
| YKR039W | *gap1Δ::KAN*, isogenic to BY4741 | Euroscarf |
| RWY176 | *pmr1Δ::NAT gap1Δ::KAN*, isogenic to BY4741 | This study |
| RWY178 | *pmr1Δ::NAT atx2Δ::KAN*, isogenic to BY4741 | This study |
| RWY056 | *pmr1Δ::NAT gdt1Δ::KAN*, isogenic to BY4741 | This study |
| NGY232 | *pmr1Δ::NAT spf1Δ::KAN*, isogenic to BY4741 | This study |
| NGY190 | *pmr1Δ::NAT ccc1Δ::KAN*, isogenic to BY4741 | R. Wellinger |
| RWY 174 | *pmr1Δ::NAT mtm1Δ::KAN*, isogenic to BY4741 | This study |
| NGY223 | *pmr1Δ::NAT pho84Δ::KAN*, isogenic to BY4741 | R. Wellinger |
| SEY6210 | *MATα leu2-3,112 ura3-52 his3-Δ200 trp-Δ901 lys2-801 suc2-Δ9 GAL* | C. Ungermann |
| CUY4517 | *OM45-GFP::HIS3*, isogenic to SEY6210 | C. Ungermann |
| RWY158 | *pmr1Δ::KAN OM45-GFP::HIS3*, isogenic to SEY6210 | This study |
| RWY189 | *smf2Δ::KAN OM45-GFP::HIS3*, isogenic to SEY6210 | This study |
| RWY190 | *pmr1Δ::NAT smf2Δ::KAN OM45-GFP::HIS3*, isogenic to SEY6210 | This study |
| NGY233 | *SMF2-GFP- CaURA3*, isogenic to BY4741 | R. Wellinger |
| MP6988 | *smf1Δ::KAN URA3::SMF1p-GFP-SMF1*, isogenic to BY4741 | This study |
| MP6994 | *smf1Δ::KAN URA3::SMF1p-GFP-SMF1 VPH1-mCherry::HIS3,* isogenic to BY4741 | This study |
| MP6998 | *SMF2-GFP- CaURA3 VPH1-mCherry::HIS3*, isogenic to BY4741 | This study |
